# A novel 3D-printed hydrogel platform for controlled delivery of BMP-9 coated calcium sulfate microparticles with co-delivery of preosteoblasts from a cell encapsulated coating layer

**DOI:** 10.64898/2026.03.18.712695

**Authors:** Janitha M. Unagolla, Ambalangodage C. Jayasuriya

**Affiliations:** Biomedical Engineering Program, Colleges of Engineering and Medicine, University of Toledo, Toledo, OH 43606, USA; Department of Orthopedic Surgery, College of Medicine and Life Sciences, 3000 Arlington Avenue, University of Toledo, Toledo, OH 43614, USA

**Keywords:** poly(ethylene glycol) dimethacrylate, 3D bioprinting, BMP-9, controlled delivery, gelatin, bone, coating layer

## Abstract

Controlled delivery of growth factors and viable cells remains a significant challenge in bone tissue engineering. In this study, a 3D-printed hydrogel scaffold system was developed for the co-delivery of bone morphogenetic protein-9 (BMP-9) and preosteoblasts to enhance bone regeneration. The system consisted of a 3D-printed base scaffold containing BMP-9-coated calcium sulfate (CaS) microparticles and a photocurable hydrogel coating layer encapsulating viable cells. The scaffold design exploited electrostatic interactions between BMP-9 and gelatin matrices by incorporating gelatin type B in the base scaffold and gelatin type A in the coating layer. Differences in the isoelectric points of these gelatin types were utilized to regulate protein binding and release. Release studies demonstrated that CaS microparticles alone exhibited rapid burst release, with nearly 80% of BMP-9 released within 24 h. Encapsulation of BMP-9 coated CaS particles in the 3D-printed scaffolds reduced the release rate, while the addition of the coating layer significantly improved sustained release, limiting BMP-9 release to approximately 50-60% by day 5. Bioactivity studies showed enhanced cell attachment in BMP-9 containing scaffolds compared with controls. Live/Dead cytotoxicity assays demonstrated high cell viability (>80%) within the coating layer over the culture period, confirming that the encapsulation and photocuring processes did not adversely affect cell survival. Cell proliferation and differentiation were further evaluated using WST-1 and alkaline phosphatase assays. The results demonstrate that electrostatic interactions governed by gelatin type selection can regulate BMP-9 release while maintaining high cell viability, providing a promising platform for growth factors and cell delivery in bone tissue engineering.

## 1. Introduction

Hydrogels have been used in many areas of tissue engineering and medicine, including oncology, immunology, cardiology, and wound healing, because of their appealing properties as drug or small molecule delivery system [1]. Hydrogels consist primarily of water (70-99%) and a crosslinked polymer network. The crosslinked hydrogel network gives hydrogels a solid-like nature, which can be tuned by changing the degree of crosslinking [2]. Hydrogels typically show a wide range of stiffness, from 0.5 kPa to 5 MPa, allowing their mechanical properties to be matched with those of different soft tissues in the human body. Also, a properly crosslinked polymer network can hinder the penetration of various proteins, preventing the premature degradation of bioactive therapeutics by inwardly diffusing enzymes. At the same time, this property supports the controlled release of bioactive therapeutics [1,3,4]. Furthermore, labile macromolecules such as recombinant proteins and monoclonal antibodies can be incorporated into hydrogels due to this intrinsic feature of crosslinked networks. Currently, more than 130 protein-based therapeutics or macromolecular drug have been approved by the Food and Drug Administration (FDA). Conventional drug administration strategies, such as intravenous and oral delivery, often require high doses and frequent repeated administration to achieve a therapeutic effect, which can lead to severe side effects and toxicity. Therefore, targeted drug delivery systems using hydrogels have gained increasing attention because of their ability to enhance therapeutic efficacy while reding dosage requirements and drug-induced toxicity [5,6].

Several hydrogel systems incorporating different liable macromolecules have been studied in the past. One of the most used hydrogel systems is calcium alginate, in which bone morphogenetic protein-2 (BMP-2)-loaded gelatin microparticles (MPs) were encapsulated as a delivery system for bone defect repair [7]. In addition, transforming growth factor–β (TGF-β) [8] and atsttrin [9], a progranulin-derived programmed protein, have also been used in calcium alginate as controlled hydrogel-based delivery system. Thermoresponsive hydrogels are also widely used in drug delivery systems. Synthetic polymers such as poly(N-isopropyl acrylamide) [10] and PCL-PEG-PCL grafted copolymers [11] have been studied by several studies for growth factor delivery. Methyl cellulose is another commonly used natural thermoresponsive polymer. An alginate/methyl cellulose thermoresponsive hydrogel system encapsulating BMP-2/9-coated chitosan MPs was developed in our lab. Excellent bioactivity was observed, as critical-size rat cranial defects healed properly without inflammatory reactions *in vivo* [12,13].

BMPs were first isolated by Marshall R. Urist in 1965 from the extract of bone [14]. BMPs are among the many growth factors that regulate cell proliferation, differentiation, and biosynthesis during bone reconstruction. To date, more than twenty BMPs have been identified, several of which play essential role in ossification, including BMP-2, BMP-4, BMP-7, and BMP-9. Among these, BMP-9 shows the strongest osteogenic potential, followed by BMP-2 and BMP-7 [15,16]. BMP-9 is also known as growth differentiation factor-2 (GDF-2), and its preprotein contains 428 amino acid residues [17]. A key characteristic of BMP-9 is that, unlike other BMPs, its signaling pathway is not inhibited by the extracellular antagonist noggin. Also, another BMP inhibitor, BMP-3, shows only a weak inhibitory effect on BMP-9 mediated osteogenesis, although it inhibits BMP-2/4/7-mediated osteogenesis pathways [18]. BMP-9 is therefore considered the most potent cytokine inducing osteogenesis among BMP family members. However, the signaling pathways underlying BMP-9-mediated osteogenesis remain poorly understood [16].

When designing a drug delivery system, the drug or macromolecule electrostatically interacts with the hydrogel or carrier to form a polyion complex [19]. These interactions between the carrier/ hydrogel and the drug typically do not dissociate simultaneously and therefore provide good stability of the polyion complex over time. Gelatin (Gel) is one of the most promising candidates for polyion complexation-based carriers due to several of its features, such as the intrinsic ability to load molecules with different charges by adjusting the pH of the solution. The isoelectric point (IEP) of the gelatin can be tailored to increase the drug loading efficiency according to the net charge of the molecule by selecting an acidic or alkaline environment. Also, drug release depends on the degradation of the hydrogel, in this case gelatin, and the degradation rate can therefore be tailored by changing the molecular weight (MW) or the degree of crosslinking [20,21].

The objective of this study was to develop a BMP-9 and cell co-delivery system for bone defect repair. In our previous study, we developed a 3D printable hydrogel system using photocurable PEGDMA. In the present study, a slightly modified version of the same 3D-printed scaffold was used to deliver both cells and BMP-9 for bone tissue engineering applications. The base scaffold was printed using the same material combination except for gelatin; instead of type A fish skin gelatin, type B bovine skin gelatin was used because the IEP of gelatin type B is more favorable for BMP-9 loading. High MW PEGDMA was synthesized in the lab using a cost-effective microwave-assisted method [22]. BMP-9 was coated onto calcium sulphate (CaS) particles in the size range of 75-106 μm to prevent the clogging of the 27-gauge nozzle during printing. The printed scaffolds were then coated with a PEGDMA hydrogel layer containing encapsulated cells. The bioactivity of BMP-9 was tested using live/dead cell assay, and cell proliferation was analyzed using the WST-1 assay. analyze the cell proliferation. BMP-9 release was evaluated using a BMP-9 ELISA kit, and cell differentiation was analyzed using an ALP assay kit.

## 2. Materials and methods

### 2.1. Materials

Poly(ethylene glycol) dimethacrylate (PEGDMA; Mn:750 Da), polyethyle glycol (PEG; MW: 6000 Da), methacrylic anhydride, diethyl ether, dichloromethane, methylcellulose (viscosity 4000 cP and 15 cP), Gelatin type A from cold-water fish skin, Gelatin type B from bovine skin, calcium sulfate (CaS) drierite, β-glycerol phosphate disodium salt pentahydrate, L-ascorbic acid, and the cell proliferation reagent WST-1 (Roche diagnostic) were purchased from Sigma Aldrich (St. Louis, MO, USA). The photoinitiator lithium phenyl-2,4,6-trimethyl benzoyl phosphinate (LAP) was purchased from Allevi (Philadelphia, PA, USA). Alpha minimum essential medium (α-MEM), fetal bovine serum (FBS), phosphate-buffered saline (PBS), Dulbecco’s phosphate-buffered saline (DPBS), and penicillin/streptomycin were purchased from Thermo Fisher Scientific through its Gibco product line. A Live/Dead cell viability/cytotoxicity kit was purchased from Invitrogen (USA). Alkaline phosphate (ALP) assay kit was purchased from Biovision (Milpitas, CA, USA). Recombinant BMP-9 and the corresponding ELISA kits containing coating, capture, and detection antibodies, along with the auxiliary kit, were purchased from R&D Systems (Minneapolis, MN).

### 2.2. Hydrogel preparation for base scaffold

LAP (0.5% w/v) was dissolved in 1x PBS solution at 65 °C, and 3% (w/v) type B bovine skin gelatin was added to the solution. After the gelatin was completely dissolved, 7% (w/v) methylcellulose (4000 cP) was added to the PBS solution. The mixture was maintained at 65 °C for 30 min to obtain a homogeneous methylcellulose solution. After 30 min, the temperature was reduced to 45 °C and PEGDMA (750 Da was added to the mixture. The resulting hydrogel precursor solution was then transferred to an ice bath to induce gelation of methylcellulose.

### 2.3. Coating of BMP-9 on the calcium sulfate microparticles

CaS flakes were ground using a mortar and pestle to obtain MPs in the size range of 75-106 μm. The particles were sieved through a 106 µm sieve, and the fraction retained on the 75 µm sieve was collected. Reconstituted BMP-9, prepared according to the manufacturer’s protocol, was then used to coat the CaS MPs. Briefly, 40 mg of MPs were coated with 4 μg of BMP-9 in 160 μl of 1% bovine serum albumin solution. The mixture was incubated at room temperature for 1 h before being mixed with the hydrogel system.

### 2.4. Printing of the base scaffolds

Scaffolds were printed using an extrusion-based Allevi2 3D bioprinter. A square mesh (5 mm ×5 mm) with four layers was designed using Autodesk Fusion 360. The CAD model was exported as stereolithography files and imported into Slic3r to generate machine-readable G-code files. The hydrogel precursor was loaded into a 10 ml syringe fitted with a 27-gauge nozzle. The printing speed was set to 6 mm/s, and the printing pressure ranged from 10-15 PSI depending on the printability of the hydrogel over time.

### 2.5. Preparation of the coating layer hydrogel

LAP (0.25% w/v) was dissolved in α-MEM at 65 °C, and 10% (w/v) PEGDMA (synthesized) was added to the solution. Subsequently, 10% (w/v) gelatin type A was added and allowed to dissolve completely. Next, 3% (w/v) methylcellulose (15 cP) was added to the solution. The mixture was maintained at 65 °C for 30 min to obtain a homogeneous MC solution and then transferred to an ice bath to induce gelation. After methylcellulose dissolution, the solution temperature was increased to 37 °C, and preosteoblasts were added at a concentration of 2 × 10⁶ cells/ml.

### 2.6. Making the coating layer around the based scaffold

Approximately 80 µL of the coating hydrogel solution was used to cover the swollen base scaffold. The coating layer was then photocured using a blue-light curing source (3M ESPE Elipar) for 120 s. Each side (top and bottom) was exposed for 60 s to ensure complete curing and prevent insufficient curing caused by blockage from the base scaffold. After curing, scaffolds with the coating layer were immersed in complete medium and incubated at 37 °C in a humidified atmosphere containing 5% CO_2_. A schematic representation of the hydrogel system consisting of the base scaffold and coating layer is shown in Fig. 1.

**Fig. 1:**
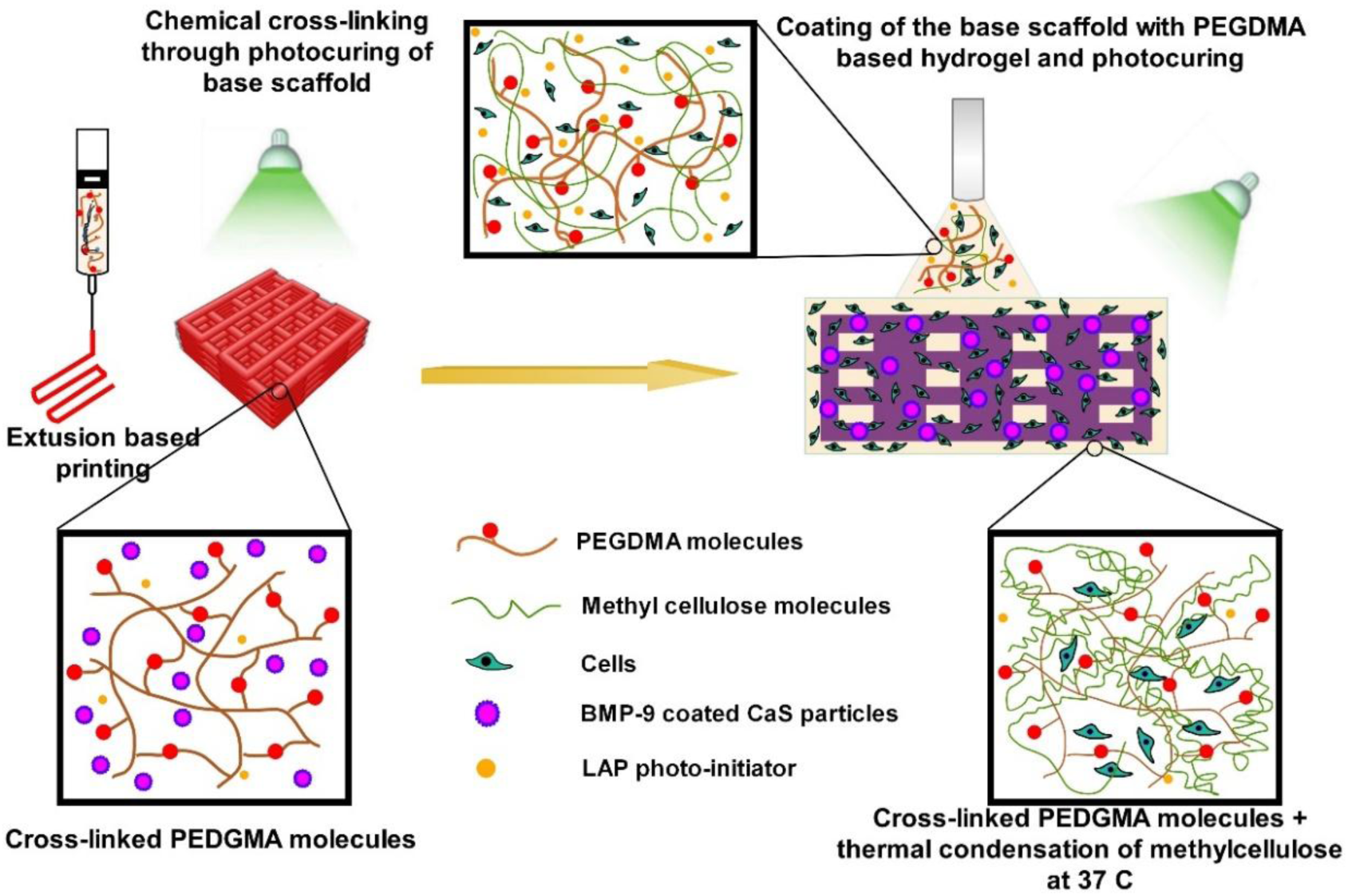
A schematic representation of the hydrogel system design consisting of a base scaffold and a coating layer.

**Fig. 1:**
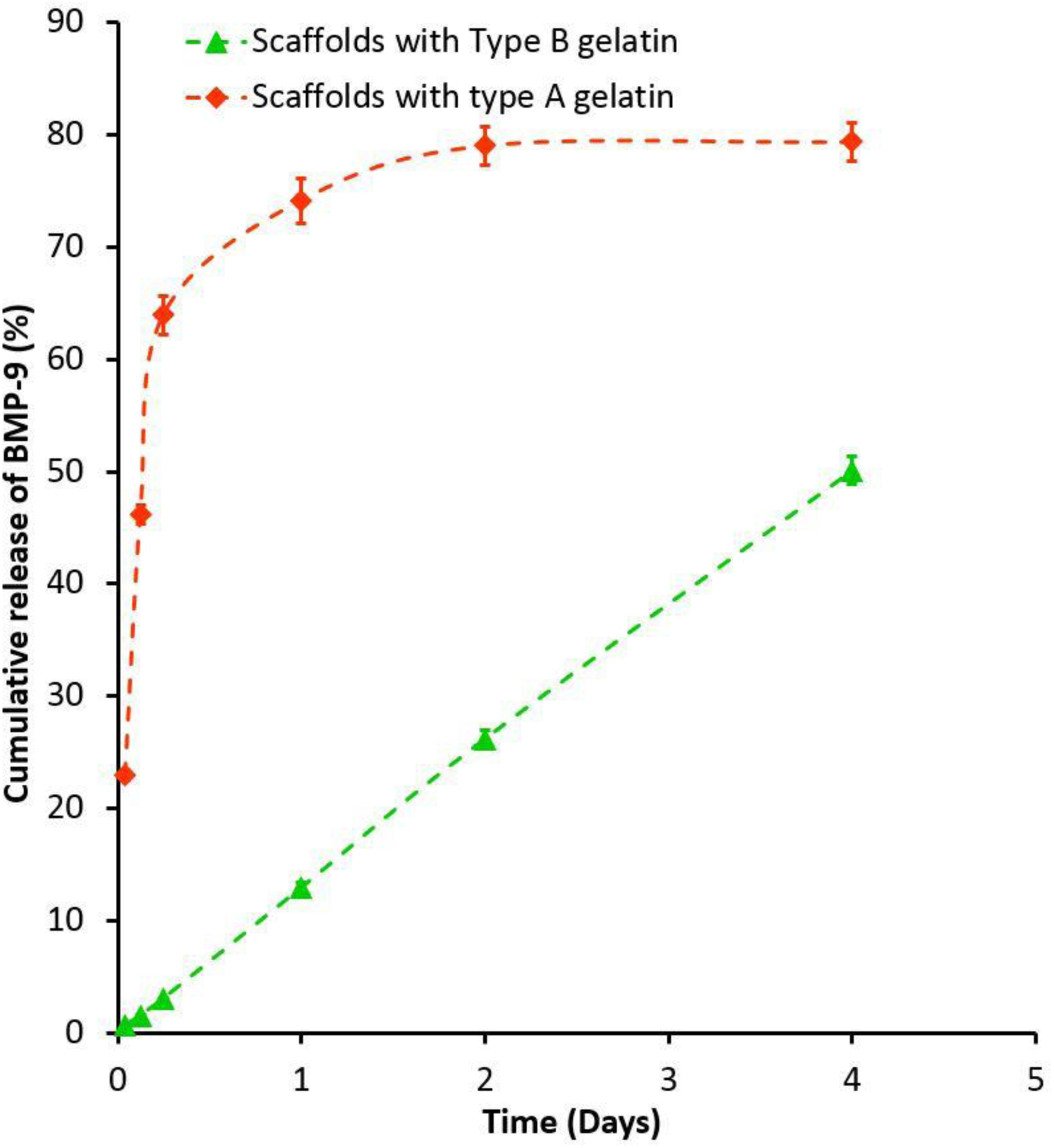
Cumulative release of BMP-9 from gelatin type A base scaffold with gelatin type A coating layer (orange); type B base scaffold with type A coating layer (light green).

### 2.7. Release study of BMP-9

BMP-9 release was studied in three experimental groups: (i) release from CaS MPs, (ii) release from uncoated scaffolds, and (iii) release from scaffolds containing the coating layer. The study was conducted at predetermined time points (1 h, 3 h, 6 h, day 1, day 2, day 4, day 7, day 10, day 14, day 18, and day 21). At each time point, 1 ml of 1x PBS was removed from the vial and replaced with fresh PBS. The collected PBS were stored at −20 °C prior to ELISA analysis. The release profile was presented as the percentage of cumulative BMP-9 release relative to the total amount initially loaded onto the MPs and scaffolds.

### 2.8. Live/Dead cytotoxicity assay

A Live/dead cytotoxicity assay was performed on the cell-encapsulated coating layer with scaffolds on days 0, 1, 3, 7, 14, and 21 according to the manufacturer’s protocol. Briefly, 5 µl of 4 mM calcein and 20 µl of 2 mM ethidium homodimer-1 were added to 10 ml of 1× DPBS to prepare the staining solution. After removing the α-MEM growth medium, scaffolds were washed with 1x PBS twice and transferred to a new 24 well plate. Then, 300 µl of staining solution was added to each scaffold and incubated at room temperature for 30 min. The top and bottom surfaces of the scaffolds were observed using fluorescence microscopy (Cytation5, BioTek Inc, USA). Images of cells attached to the well plate beneath the scaffolds were also captured to evaluate cell migration and proliferation. Images were processed and merged using Gen 3.03 software. Cell viability was quantified using ImageJ according to the manufacturer’s protocol. Five images per time point were used for quantification.

### 2.9. Cell proliferation assay

Cell proliferation was quantified using the WST-1 assay, which measures cellular metabolic activity that correlates with the number of viable cells through the formation of a formazan dye.

The assay was performed on days 1, 3, 7, and 14. At each time point, α-MEM medium was removed, and scaffolds were washed with 1× PBS. Fresh α-MEM medium containing 10% (v/v) WST-1 reagent was added to the scaffolds after transferring them to new wells. Blank samples containing scaffolds without cells were also prepared. The plates were shaken at 300 rpm for 2 min to ensure homogeneous mixing and then incubated for 4 h at 37 °C and 5% CO₂. After incubation, 100 µl from each well was transferred to a 96-well plate, and absorbance was measured at 440 nm using a spectrophotometer (SpectraMax 190). Background absorbance from blank samples was subtracted from each experimental value.

### 2.10. Alkaline phosphatase (ALP) cell differentiation assay

Cellular ALP activity was determined by quantifying p-nitrophenol, which is produced by dephosphorylation of p-nitrophenyl phosphate (pNPP) in the presence of ALP. For the assay, scaffolds containing the cell-encapsulated coating layer were washed thoroughly with 1× PBS and transferred to microcentrifuge tubes. Next, 300 µl of lysis buffer was added and the samples were vortexed for 1 min to lyse the cells. Prior to vortexing, the coating layer and scaffold were crushed into small pieces to facilitate cell lysis. The lysates were kept on ice for approximately 20 min and then centrifuged at 13,000 g for 3 min. The supernatant was collected and centrifuged again. The final supernatant was mixed with the pNPP substrate, and the assay was performed according to the kit protocol. Absorbance was measured using the SpectraMax 190 microplate reader, and ALP activity was calculated based on the standard curve obtained using the ALP enzyme provided in the kit. The measured ALP activity was normalized to the total protein content in each sample. Total protein concentration was determined using the Coomassie Plus protein assay kit (Thermo Scientific) according to the manufacturer’s instructions.

## 3. Results

### 3.1. BMP-9 release study

First, the proposed hypothesis of using gelatin type A and type B in the coating layer and base scaffold was evaluated using the BMP-coated CaS particles encapsulated within the scaffolds. The release data showed the proposed hypothesis was valid, as burst release was observed in the scaffold system where gelatin type A was used for both the base scaffold and the coating layer (Fig. 2). Therefore, by carefully selecting the IEP of the materials, the release of the biological molecules can be appropriately controlled.

**Fig. 2:**
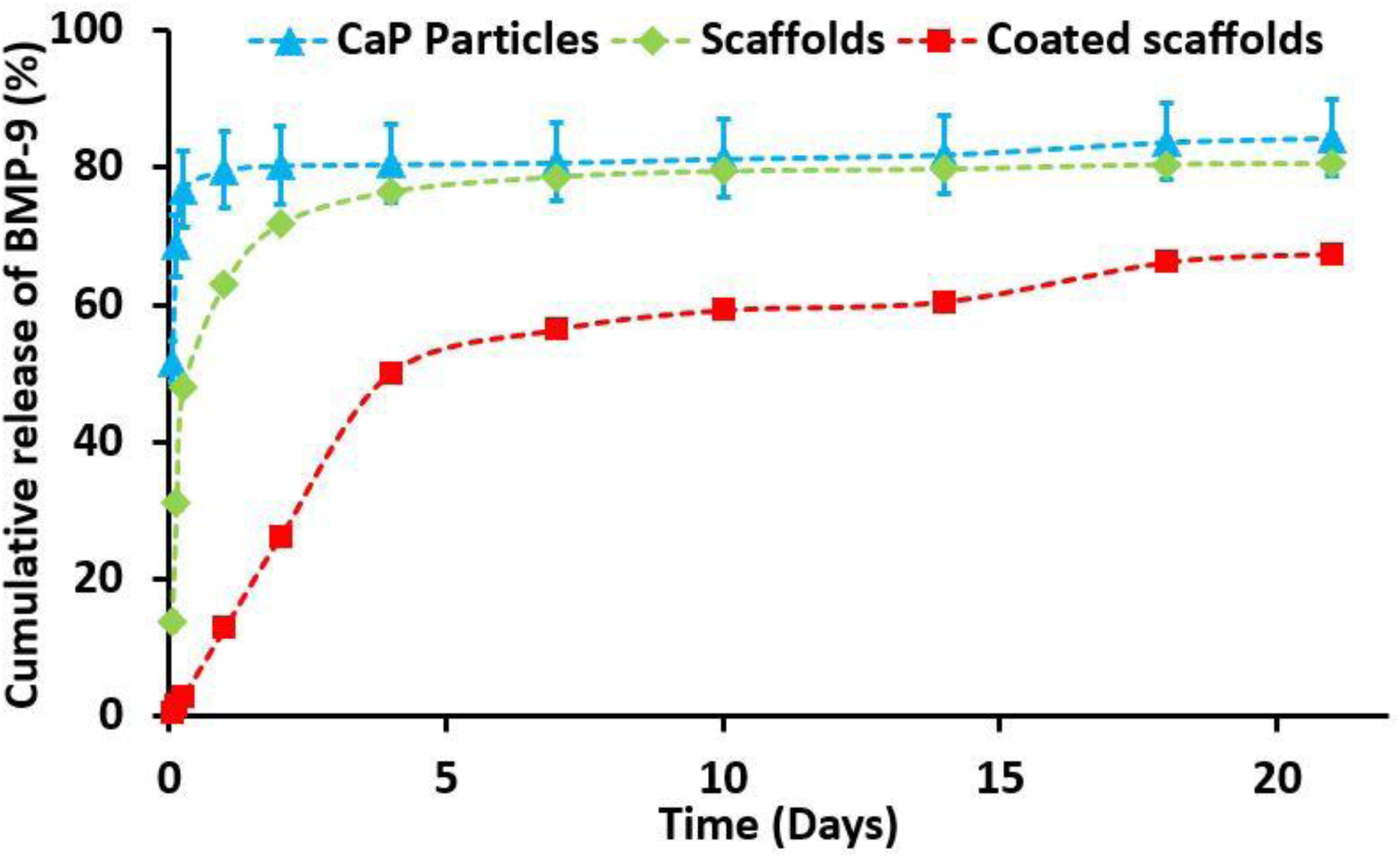
Cumulative release of BMP-9 over 21 days, n=3.

Figure 3 shows the cumulative release of BMP-9 from CaS particles, CaS (with BMP-9) encapsulated 3D-printed scaffolds, and BMP-9 from coated scaffolds. As expected, CaS MPs showed a burst release, and within one day, nearly 80% of the BMP-9 was released to the PBS solution. Scaffolds without a coating layer also showed a higher release rate of BMP-9, with almost 89% of the BMP-9 released by day 5, compared to the 80% release from CaS MPs on day 1. The coating layer played a major role in controlling BMP-9 release, as only 50–60% was released by day 5. These results confirm the validity of the proposed hypothesis.

**Fig. 3:**
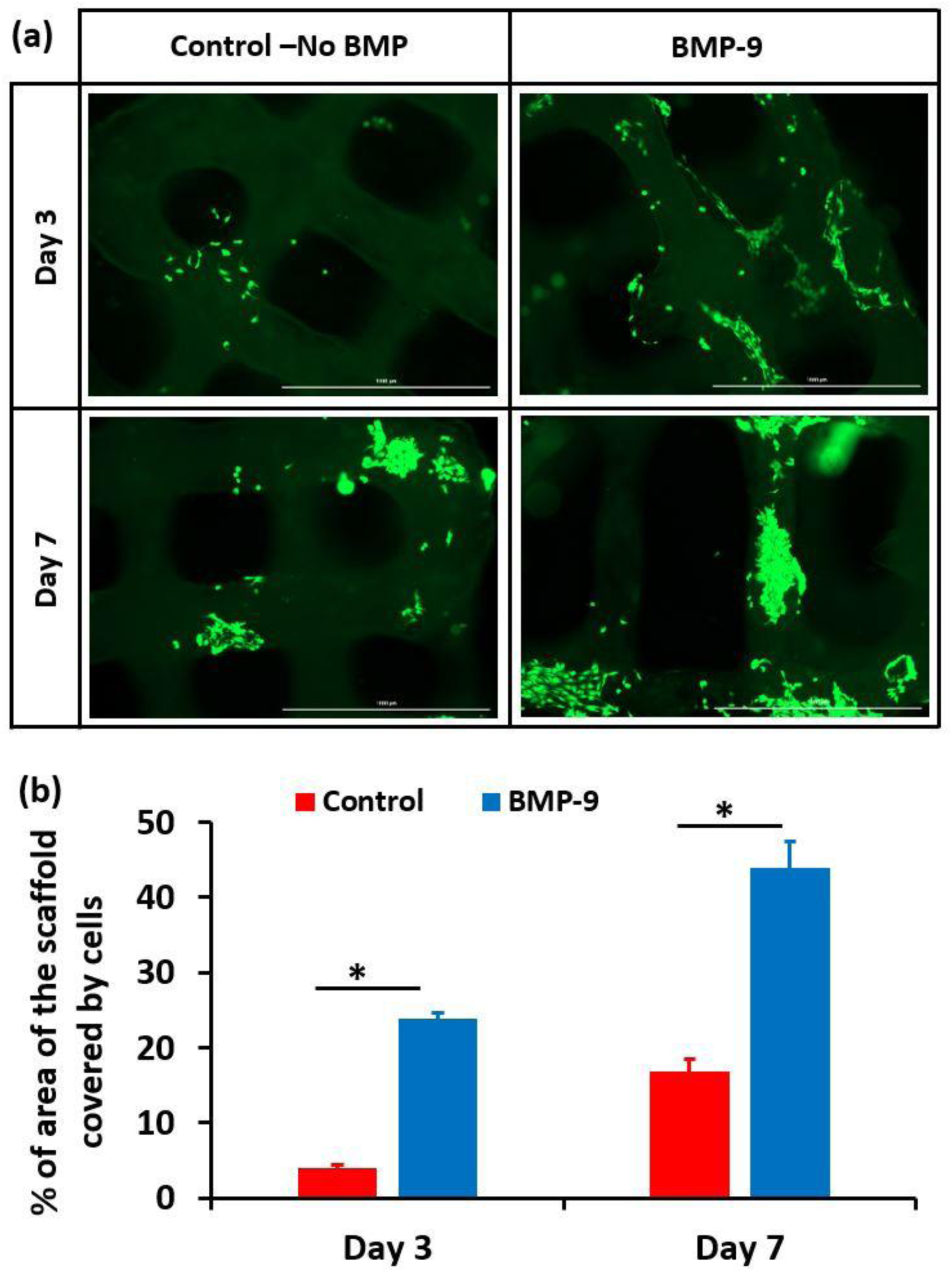
Bioactivity of BMP-9: (a) Live cell assay images at days 3 and 7 for BMP-9 coated CaS MPs encapsulated scaffolds and control scaffolds without BMP-9; (b) Quantification of live cell images using Image J software; n=5, * indicates the significance p<0.05.

### 3.2. Bioactivity of the BMP-9

The bioactivity induced by BMP-9 was evaluated using the base scaffolds by examining cell attachment. Fig. 4 shows a significantly higher level of cell attachment in the BMP-9-encapsulated scaffolds compared to scaffolds without BMP-9. A statistically significant difference was observed between the two groups, suggesting improved scaffold bioactivity due to BMP-9.

**Fig. 4:**
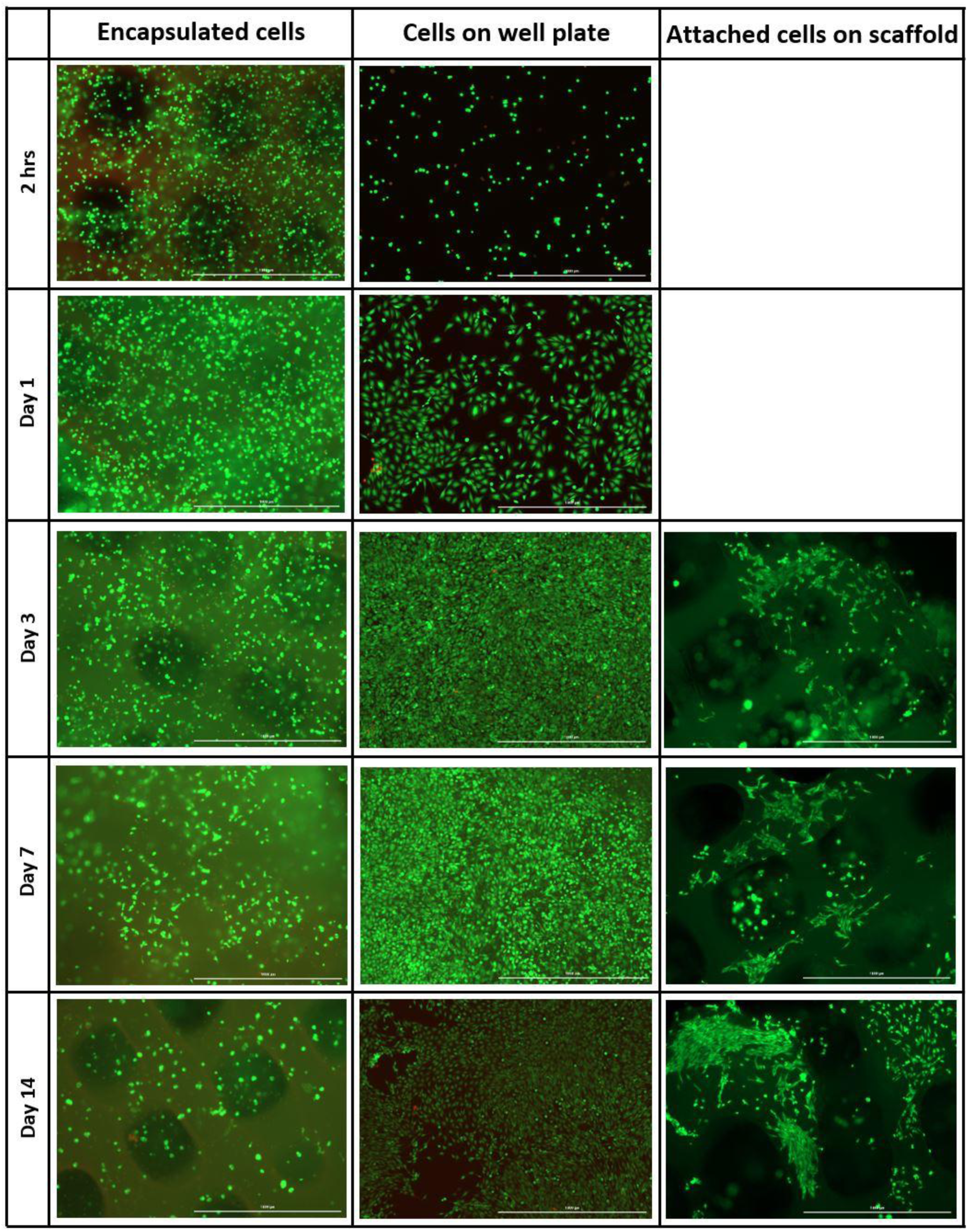
Live/Dead cell assay images of the base scaffold coated PEGDMA coating layer; cells released from the coating layer and proliferated on the well plate and after removing the coating layer, cells attached to the base scaffolds; green-calceing, red-edithelium homodimer.

### 3.3. Cell viability of the coating layer

The LD assay images are shown in Fig. 5. On day 0 or after 2 h of cell encapsulation in the coating layer, very high cell viability was observed. The cells had not escaped from the coating layer, and few cells were visible at the bottom of the plate. On day 1, the coating layer contained more viable cells, as shown in Fig. 5, but some cells had escaped from the coating layer and attached to the well plate. This suggests that the coating layer is not toxic to the encapsulated cells and that the cells remain healthy enough to proliferate after the mixing and photocuring steps.

**Fig. 5:**
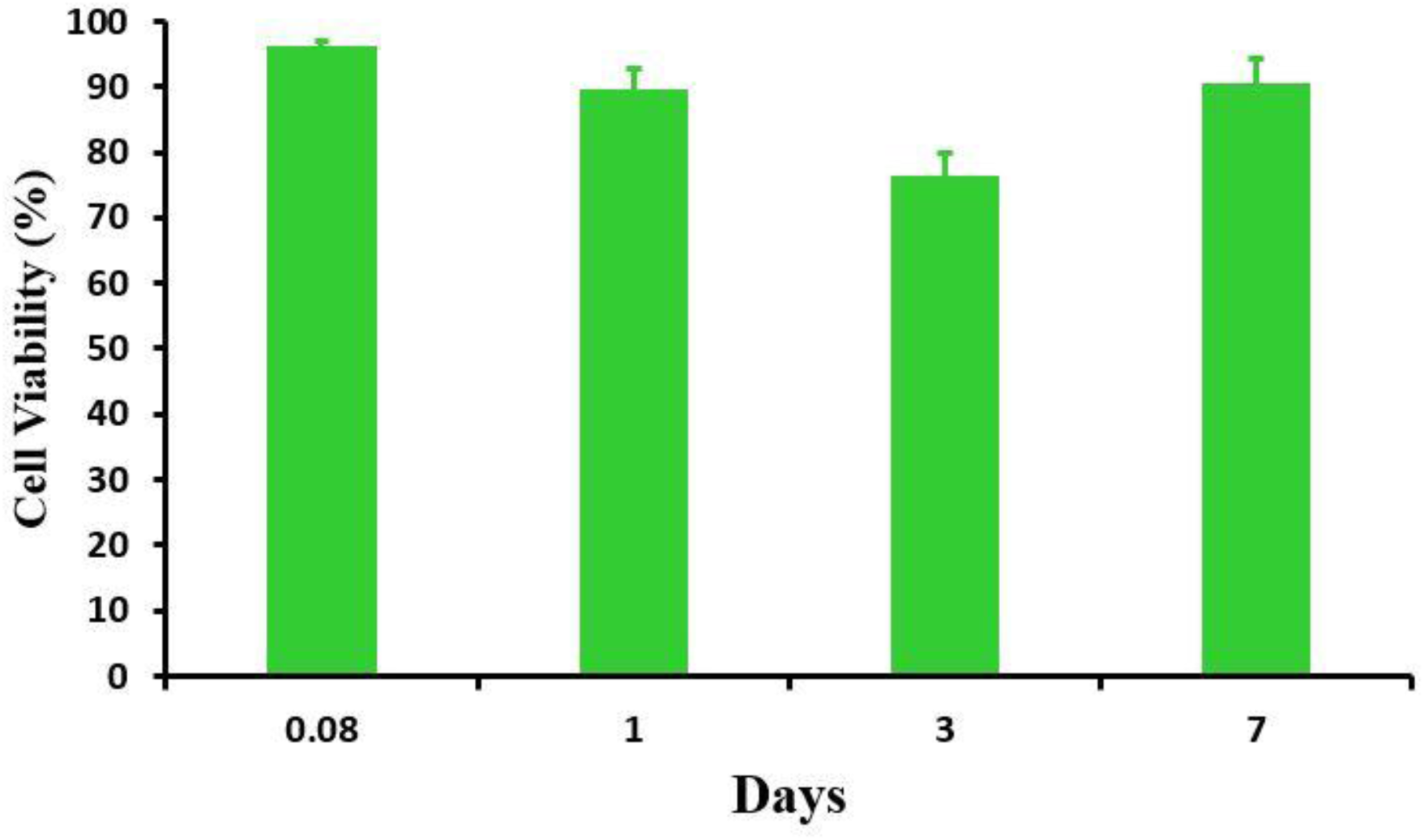
Cell viability inside the coating layer as a percentage of total live and dead cells using Image J software; n=5.

A similar behavior was observed on day 3, when the well plates were completely covered with the cells, and more cells had been released from the coating layer. In addition, the escaped cells were attached to the base scaffolds. On days 7 and 14, more cells escaped from the coating layer, and these cells attached to and proliferated on the well plate. However, for day 14, the scaffolds had been transferred to a new well plate on day 7 due to excessive cell growth, which resulted in lower cell proliferation on day 14 well plate, as shown in Fig. 5. Additionally, more cells attached to the base scaffold, and the coating layer began to degrade.

The quantitative cell viability is shown in Fig. 6. As described earlier, the coating layer exhibits excellent cell viability. There was a gradual decline in cell viability up to day 3, followed by a gradual increase. The lowest cell viability (80%) was observed on day 3. By day 7, more cells had escaped from the coating layer, however, the remaining cells within the coating layer were viable, contributing to an overall increase in cell viability.

**Fig. 6:**
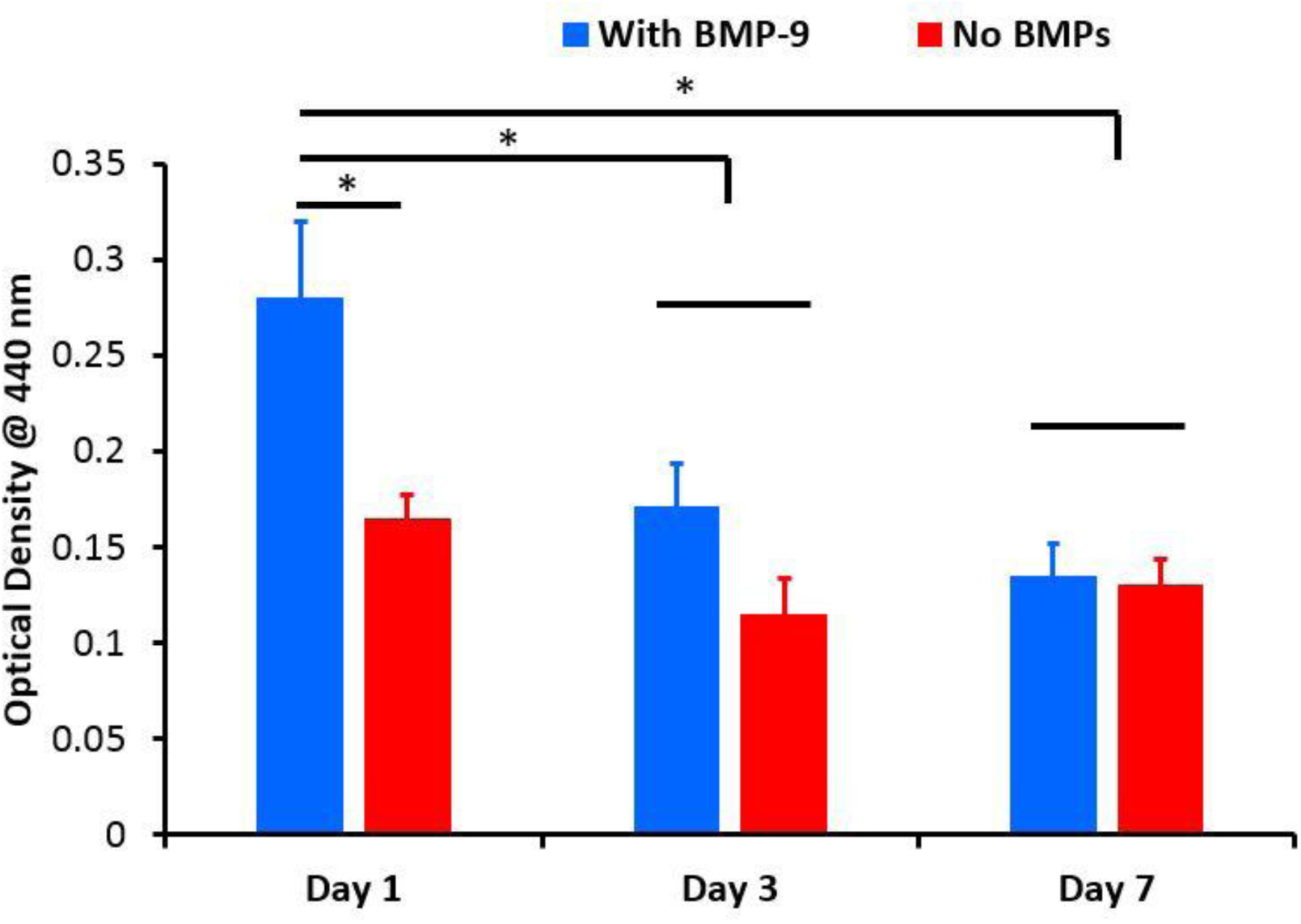
WST-1 cell proliferation assay results; the OD values are given since it is proportional to the number of cells; n=3, * indicates the significance p<0.05.

### 3.4. Cell proliferation and differentiation of encapsulated cells with BMP-9

Figure 7 shows the WST-1 cell proliferation assay results of BMP-9 encapsulated scaffolds with a coating layer and scaffolds without BMP-9 but with the coating layer. As previously observed in the Live/Dead assay images, a similar trend was observed. Day 1 showed the highest optical density value, suggesting a higher number of cells in the coating layer. There was a significant difference between the BMP-9 containing sample and the without BMP-9 control. However, similar to the Live/Dead assay results, optical density values gradually decreased over time, and no significant differences were observed between groups on days 3 and 7, suggesting the release of cells and their attachment to the base scaffolds.

**Fig. 7:**
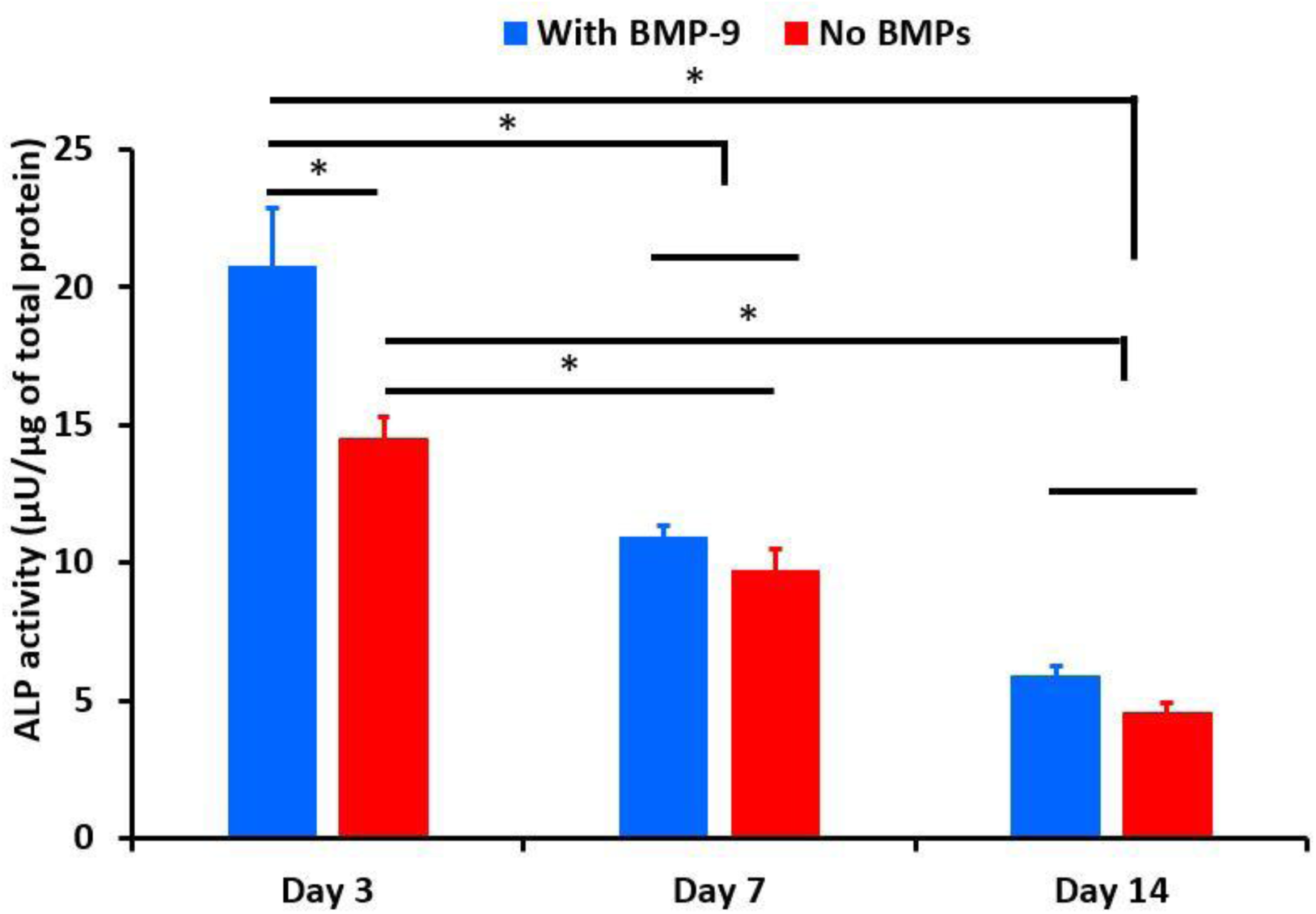
ALP activity of the cells encapsulated in the coating layer; n=3, * indicates the significance p<0.05.

Figure 8 shows the ALP activity assay results for BMP-9 containing and non-BMP-9 containing coated scaffolds. Again, a similar trend to the WST-1 assay was observed. There was a significant difference between the BMP-9 containing scaffolds and control, as expected. However, the gradual reduction in ALP activity can be attributed to the decrease in the number of encapsulated cells within the coating layer over time.

**Fig. 8:**
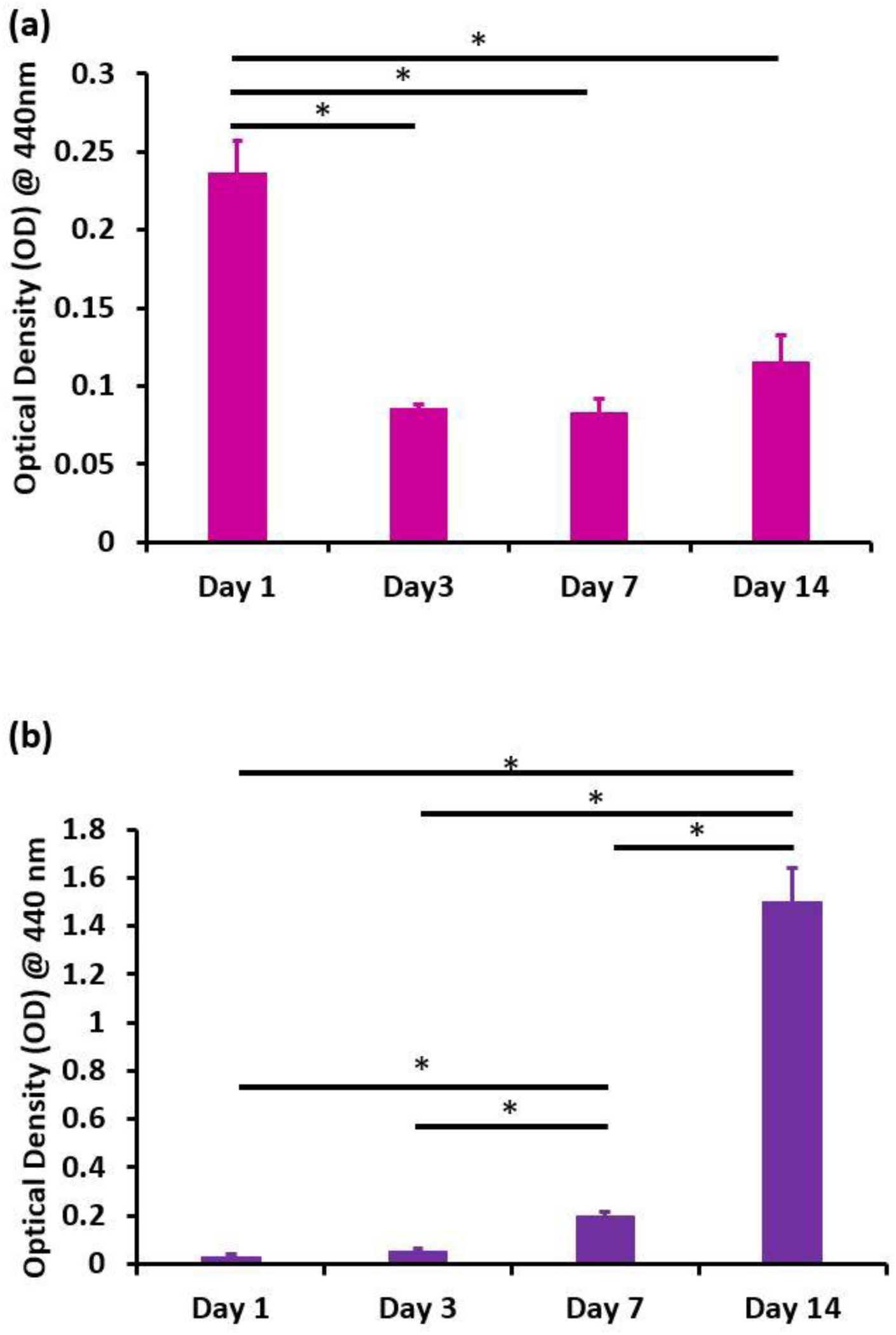
WST-1 assay experiments on cells encapsulated on the coating layer; (a) cells in the coating layer; (b) cells proliferate on the well plate.

Since the results showed reduced cell proliferation and ALP activity over time in the coating layer, the experiments were repeated to further clarify these findings. Figure 9 shows the WST-1 assay results for the cell-encapsulated coating layer, however, BMP-9 was not used in this experiment. Consistent with previous results, there was a decrease in the number of cells on day 3, likely due to cell escape from the coating layer. After day 3, the number of cells in the coating remained relatively constant on day 7, suggesting cell attachment to the base scaffold (Fig. 9(b)). By day 14, the number of cells increased as expected due to continued cell attachment and proliferation. In contrast, the number of viable cells in the well plate continuously increased, as shown in Fig. 9(b). These results are consistent with the Live/Dead assay results shown previously in Fig. 5.

**Fig. 9:**
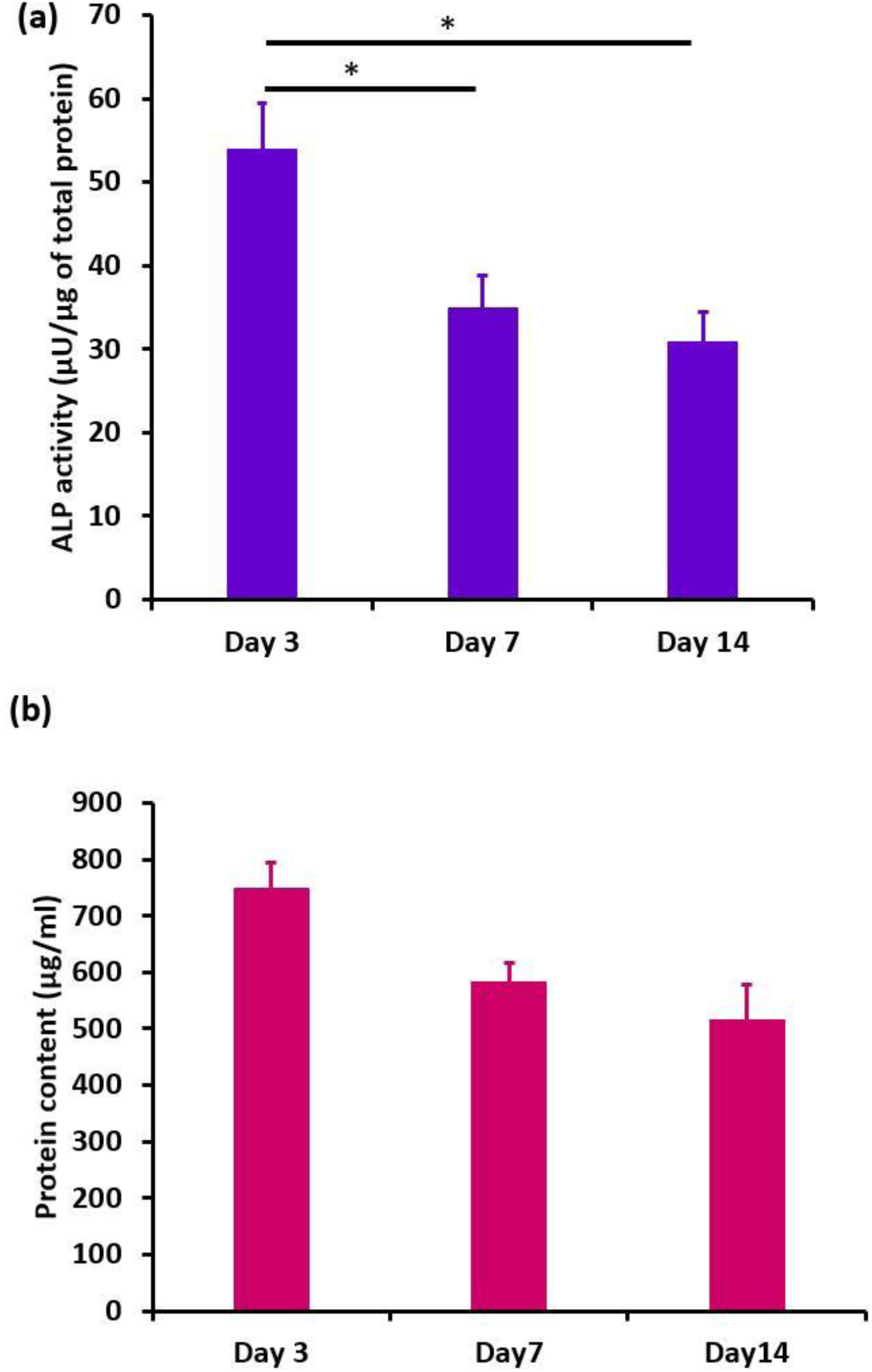
(a) ALP activity of the cells differentiated inside the coating layer that normalized with the total protein content (b) total protein content of the cells inside the coating layer; n=3, * indicates the significance p<0.05.

Figure 10(a) shows the ALP activity of the cells encapsulated within the coating layer. A similar trend was observed before: after day 3, there was no significant difference between days 7 and 14, suggesting the cell proliferation on the base scaffold. In addition, ALP activity was normalized to the total protein content of each coating layer. The total protein content is shown in Fig. 10(b), and no significant differences were observed between the groups, further supporting the WST-1 assay results on days 3, 7, and 14.

## 4. Discussion

The encapsulation of drugs, peptides, and proteins, and their controlled release remain significant challenges in the development drug delivery systems. In this study, BMP-9 was coated on CaS MPs as a delivery vehicle within a hydrogel system. Preliminary studies showed that directly mixing the protein with the hydrogel resulted in burst release, with nearly 90% of the encapsulated BMP-9 released on day 1.

CaS MPs are widely used for drug and protein delivery [23,24]. In addition to this function, CaS exhibits excellent osteoconductive properties. *In vivo*, in the presence of bone, CaS becomes osteogenic and is completely absorbed without significant inflammatory response [25,26]. Therefore, CaS serves both as a delivery vehicle and as an osteogenic enhancer in bone tissue engineering applications.

The hydrophilic nature of BMP-9 promotes burst release from CaS MPs. To address this, a hydrogel system was designed to reduce this effect. The binding of the BMPs depends on electrostatic interactions between the protein and the hydrogel. Several factors influence this interaction, including ionic strength, pH of the solution, IEP, and polymer composition, as release is predominately governed by electrostatic mechanisms [27]. This type of mechanism was proposed for the first time for BMPs.

The IEP of gelatin type A is approximately pH 9, while gelatin type B has an IEP around pH 5. BMPs generally have an IEP in the range of pH 7.7-9. At physiological pH (7.4), BMP-9 carries a positive charge. Gelatin type A is also positively charged, resulting in electrostatic repulsion, whereas gelatin type B is negatively charged, leading to electrostatic attraction. In this system, BMP-9 coated MPs were encapsulated in a gelatin type B matrix, promoting attraction, while the coating layer consisted of gelatin type A, inducing repulsion. These combined interactions regulate the release of BMP-9, as illustrated in Fig. 3. This approach may be extended to other hydrogel systems and bioactive molecules.

BMP-9 is one of the least studied growth factors in the TGF-β superfamily. A common strategy for inducing endogenous BMP-9 expression involves transfecting stem cells using viral vectors and seeding those cells on the scaffolds for bone regeneration [28–30]. Previous work in our lab demonstrated significant osteogenesis using recombinant BMP-9 (rhBMP-9) delivered via a thermoresponsive hydrogel system in a rat model [12,13]. The coating layer-based system proposed here supports sustained BMP-9 release and can be adapted to complex defect geometries using 3D-printed scaffolds.

The coating layer was designed to be fragile to support initial cell viability and facilitate in vivo cell delivery. PEG (6 kDa) was selected as an optimal material. High initial cell viability and early cell release (within one day) indicate minimal stress during encapsulation and photocuring. Released cells proliferated in the well plate, confirming cell health (Fig. 6). Cell release from photocurable hydrogels has been reported in previous studies. [31,32].

WST-1 and ALP assays further supported these observations. Cells released from the coating layer proliferated and eventually attached to the base scaffold as the coating degraded. In the ALP assay, only encapsulated cells were analyzed. Therefore, the decrease in ALP activity from day 3 to day 7 reflects the reduction of cells within the coating layer, while the similar ALP activity from day 7 to day 14 indicates proliferation on the base scaffold.

*In vivo*, cells released from the coating layer are expected to proliferate at the defect site and differentiate into osteoblasts, enhancing osteogenesis in combination with BMP-9 signaling. Also, this system can be further investigated in different proteins and peptides. By appropriately selecting the IEP, this system may also be adapted for the delivery of other proteins, peptides, and small molecules by tuning electrostatic interactions.

## 5. Conclusion

A 3D-printed hydrogel scaffold system capable of co-delivering BMP-9 and viable cells was successfully developed in this study. The system utilized BMP-9 coated CaS MPs incorporated into a gelatin-based hydrogel scaffold with a photocurable coating layer. By strategically selecting gelatin type B for the base scaffold and gelatin type A for the coating layer, electrostatic interactions based on differences in isoelectric points were exploited to regulate protein release.

The results demonstrated that CaS MPs alone exhibited rapid burst release of BMP-9, whereas encapsulation within the hydrogel scaffold significantly slowed the release. The addition of the coating layer further improved release control, resulting in sustained BMP-9 delivery over several days. Bioactivity studies confirmed that BMP-9 enhanced cell attachment and maintained biological activity within the scaffold system. The coating layer supported high cell viability and allowed gradual cell release and proliferation on the scaffold surface.

These findings suggest that controlling electrostatic interactions within hydrogel systems provides an effective strategy for regulating growth factor release. The proposed scaffold-coating layer platform offers a versatile approach for the co-delivery of cells and bioactive molecules and may be applied to a wide range of growth factors, peptides, and therapeutic proteins for bone tissue engineering and regenerative medicine.

## Acknowledgment

We would like to acknowledge National Institutes of Health (NIH) grant number R01DE023356.

## Data Availability

The data used to evaluate the findings of this study are present in the paper.

